# Feeding dihydroquercetin and vitamin E to broiler chickens reared at standard and high ambient temperatures

**DOI:** 10.1101/2020.05.19.104398

**Authors:** V. Pirgozliev, C. Westbrook, S. Woods, S.C. Mansbridge, S.P. Rose, I.M. Whiting, D. Yovchev, A.G. Atanasov, K. Kljak, G.P. Staykova, S. Ivanova, M.R. Karagecili, F. Karadas, J.H. Stringhini

**Affiliations:** The National Institute of Poultry Husbandry, Harper Adams University, Shropshire, UK; Faculty of Veterinary Medicine, Trakia University, 6000 Stara Zagora, Bulgaria; Ludwig Boltzmann Institute for Digital Health and Patient Safety, Medical University of Vienna, 1090 Vienna, Austria; Institute of Genetics and Animal Biotechnology of the Polish Academy of Sciences, 05-552 Magdalenka, Poland; Institute of Neurobiology, Bulgarian Academy of Sciences, 1113 Sofia, Bulgaria; Department of Pharmacognosy, University of Vienna, 1090 Vienna, Austria; Faculty of Agriculture, University of Zagreb, Croatia; Agricultural Institute, 9700 Shumen, Bulgaria; Agricultural Academy, 1373 Sofia, Bulgaria; Department of Animal Science, Yuzuncu Yil University, Van, Turkey; Universidade Federal de Goias, Goiania, Brazil

**Keywords:** broilers, dihydroquercetin (DHQ), vitamin E, growth performance, GSH-Px, ambient temperature

## Abstract

The use of natural antioxidants, in particular polyphenols such as dihydroquercetin (DHQ), in animal nutrition have recently increased in popularity. This may partly be due to the risk of increased incidences of heat stress associated with raising livestock in warmer ambient temperatures, facilitated by global warming, reducing antioxidant capacity. The current research demonstrates the effect of dietary DHQ, vitamin E and standard or high ambient temperatures on growth performance, energy and nutrient metabolism, gastrointestinal tract development (GIT), jejunal villus morphometry and antioxidant status in broiler chickens. Each of the four experimental diets were fed to 16 pens of five birds, which were allocated to four rooms (four pens in each room). The temperature in two rooms was maintained at a constant 35 °C (high temperature; HT), and the temperature in the other two rooms was gradually reduced from 27 °C at 7d of age to 22 °C at 20d of age (standard temperature; ST). Rearing birds at HT reduced: feed intake, weight gain, weight of small intestine, total GIT, liver, spleen, heart, villus height, villus surface area and lowered blood glutationperoxidase (GSH-Px). Dietary DHQ increased blood GSH-Px and total antioxidant status, increased heart weight and reduced caecal size. When fed separately, DHQ and vitamin E improved hepatic vitamin E concentration. Feeding vitamin E increased spleen and liver weights. When fed together, DHQ and vitamin E reduced villus height, villus height to crypt depth ratio and villus surface area. Temperature and antioxidants did not affect energy and nutrient metabolism. There were no effects of dietary antioxidants on growth performance of broiler chickens and there were no mortalities. At present it is unclear if feeding antioxidants (in particular DHQ) at different levels, using different dietary formulations, and rearing birds under a range of environmental conditions may be effective at enhancing production performance and bird health in hot ambient climates.

## 1. INTRODUCTION

It is well documented that high ambient temperatures adversely affect production and physiology in poultry (Niu et al., 2009; Quinteiro-Filho et al., 2010; Woods et al., 2020a). Heat stress is an environmental factor which leads to oxidative stress, namely the disruption of the equilibrium between antioxidants and reactive oxygen species (ROS) (Gu et al. 2012). Since the global climate is changing and getting warmer, the rise in temperature is an increasingly important consideration for poultry producers (Nawab et al., 2018). To reduce the impact of high temperatures, producers in hot climates typically use cooling and ventilation systems which increase production costs and are only applicable in intensive production systems (Al-Murrani et al., 1997). However, the use of free-range rearing systems in broiler production is increasing, thus research into different approaches to alleviate the impact of heat stress is needed.

The use of natural antioxidants, in particular polyphenols, in food and nutrition has recently gained increased popularity (Surai, 2014; Iskender et al., 2017). Dihydroquercetin (DHQ), also known as taxifolin, is a flavonoid, a major sub-group representing plant polyphenols, commonly found in onions, milk thistle, and various conifers (Weidmann, 2012). Dihydroquercetin has been widely applied as an antioxidant for the surface treatment of fresh meat and fish (Kamboh et al., 2019). Dihydroquercetin has also been incorporated in animal diets in order to enhance meat quality and oxidative stability of pork (Vlahova-Vangelova et al., 2020). An extensive review by Fomichev et al. (2017) reported an enhancement in growth performance of poultry and pigs when fed DHQ supplemented diets, although the responses were more noticeable during summer months. Balev et al. (2015) and more recently Pirgozliev et al. (2019a) did not find significant differences in growth performance or physiological variables of fully-grown broilers fed DHQ, when reared under industry conditions. It has been suggested, however, that where reported improvements in production variables have been noted in the literature, these may be observed when animals are exposed to heat stress (Fomichev et al. 2017). Rearing animals at temperatures approaching or exceeding their thermal comfort zone, i.e. during summer, may be a reason for depleting levels of tissue antioxidants; thus, the antioxidant status of animals may be enhanced by dietary DHQ supplementation, since it is a natural flavonoid with recognised antioxidant properties (Surai 2014, 2016). However, there are no reported studies comparing the response to DHQ of broilers reared under standard and high ambient temperatures. In addition, there are no comparisons between the effectiveness of DHQ and other well recognised antioxidants, e.g. vitamin E, on their impact (and interactions) on growth performance and antioxidant capacity of poultry at different rearing temperatures. Dietary inclusion of supplementary antioxidants, including polyphenols and vitamin E, have been shown to reduce the adverse impact of high temperature and improve antioxidant status and growth performance of poultry (Fomichev et al. 2017; Mazur-Kuśnirek et al., 2019).

The primary objectives of this experiment were to study the impact of dietary DHQ and vitamin E on growth performance variables, dietary N-corrected apparent metabolisable energy (AMEn), dry matter (DMR) and nitrogen retention (NR) coefficients when fed to broiler chickens from 7 to 28 days of age, reared at industry recommended and high ambient temperatures. In addition, secondary objectives were to examine the impact of experimental diets and ambient temperatures on gastrointestinal tract (GIT) and internal organ development, and jejunal villus morphometry. Finally, an evaluation of the influence of antioxidants and ambient temperatures on hepatic vitamin E, glutathione peroxidase (GSH-Px) in blood, total antioxidant status (TAS), heterophil to lymphocyte (H:L) ratio and packed cell volume (PCV) was conducted.

## 2. MATERIALS AND METHODS

### 2.1. Experimental diets

A wheat-soy-based basal grower diet formulated to meet breeder’s recommendations (Aviagen Ltd., Edinburgh, UK) (Table 1) was mixed for the experiment. The diet was supplied with 5 g/kg of TiO_2_ as an indigestible marker. The basal diet was then split into four batches that had 1.) no additive (control diet; C); 2.) C + 0.5 g/kg extract of Siberian Larch (*Larix sibirica*) (JSC NPF Flavit, IBI RAS, Pushchino city, Moscow region, Russian Federation 142290). According to the supplier, this extract contains over 85 % pure DHQ (DHQ diet); 3.) C + 0.3 g/kg vitamin E (Merck KGaA, Darmstadt, Germany) (vit E diet); 4.) C + 0.5 g/kg extract of Siberian Larch (*Larix sibirica*) + 0.3 g/kg vitamin E (DHQ + vit E diet).

**Table 1.** Ingredient composition (g/kg ‘as fed’) and nutritional analysis of the basal diet for broiler chickens

| Ingredients | g/kg |
| --- | --- |
| Barley | 79 |
| Wheat | 550 |
| Soya bean meal | 230 |
| Soya full fat | 50 |
| L Lysine HCL | 3 |
| DL Methionine | 3.5 |
| L Threonine | 1.5 |
| Soya oil | 45 |
| Limestone Trucal 52 | 12.5 |
| Monocalcium phopshate | 12.5 |
| Salt | 2.5 |
| Sodium bicarbonate | 1.5 |
| Premix<sup>1</sup> | 4 |
| Titanium Dioxide | 5 |
| Calculated values (as fed) |  |
| Crude protein (Nx6.25) | 201 |
| Crude oil | 68 |
| ME, MJ/kg | 12.99 |
| Calcium | 9.3 |
| Av Phosphorus | 4.2 |
| Av Lysine | 11.8 |
| Methionine + Cysteine | 9.4 |
| Determined values |  |
| DM | 894 |
| GE MJ/kg | 17.43 |
| Crude protein (N x 6.25) | 194 |
| Crude oil | 69 |
| Vitamin E (µg/g)<sup>2</sup> | 43.86 |

<sup>1</sup>The Vitamin and mineral premix contained vitamins and trace elements to meet the requirements specified by NRC (1994) except vitamin E. There was no vitamin E supplemented to the control diet. The premix provided (units/kg diet): retinol 3600 µg, cholecalciferol 125 µg, menadione 3 mg, thiamine 2 mg, riboflavin 7 mg, pyridoxine 5 mg, cobalamin 15 µg, nicotinic acid 50 mg, pantotenic acid 15 mg, folic acid 1 mg, biotin 200 µg, iron 80 mg, copper 10 mg, manganese 100 mg, cobalt 0.5 mg, zinc 80 mg, iodine 1 mg, selenium 0.2 mg and molybdenum 0.5 mg.
<sup>2</sup>The determined values of vitamin E (µg/g) for diets 2, 3 and 4 are 54.31, 87.51 and 83.49, respectively.

### 2.2. Husbandry and sample collection

The experiment was conducted at the National Institute of Poultry Husbandry and approved by the Research Ethics Committee of Harper Adams University, UK. A total of 340 day-old male Ross 308 broilers were obtained from a commercial hatchery (Cyril Bason Ltd, Craven Arms, UK), allocated to a single floor pen and offered a proprietary wheat-based broiler starter feed formulated to meet Ross 308 nutrient requirements (Aviagen Ltd., Edinburgh, UK). At 7d age, 320 of the birds, excluding ill and malformed, were allocated at random to the four experimental diets. Each diet was fed to 16 pens (containing five birds) in total, which were allocated to four rooms, four pens in each room. Each of the pens had a solid floor and were equipped with an individual feeder and drinker. Feed and water were offered *ad libitum* to birds throughout the experiment. The temperature in two of the rooms was maintained at a constant 35 °C (HT), and the temperature in the other two rooms was gradually reduced from 27 °C at 7d age to 22 °C at 20d age (following breeder’s recommendations; ST). A standard lighting programme for broilers was used, decreasing the light:dark ratio from 23h:1h from day old to 18h:6h at 7d of age, which was maintained until the end of the study. The well-being of the birds was checked regularly every day.

Birds and feed were weighed on days 7 and 28 in order to determine average daily feed intake (FI), average daily weight gain (WG) and to calculate the feed conversion ratio (FCR) on a pen basis. For the last three days of the study, from day 18 to day 21, the solid floor of each pen was replaced with a wire mesh. During this period excreta were collected each day, stored in a fridge (∼5 °C), and a well-homogenised representative subsample was dried at 60 °C and then milled through a 0.75 mm screen.

At the end of the study, one bird per pen, selected at random, was electrically stunned and blood was obtained in heparin coated tubes from the jugular vein. The development of the GIT from the same birds was determined. The proventriculus and gizzard (PG), duodenum, pancreas, jejunum, ileum, caeca, liver, spleen and the heart were immediately collected and weighed. The liver was freeze dried and stored at minus 80 °C before being analysed for vitamin E content. Approximately 5 cm of the middle part of the jejunum, between the point of bile duct entry and Meckel’s diverticulum, of one of the birds was sampled and stored in 10 % neutral-buffered formalin before further processing.

### 2.3. Laboratory Analysis

Dry matter (DM) in feed and excreta samples was determined by drying of samples in a forced draft oven at 105 °C to a constant weight (AOAC 2000; method 934.01). Crude protein (6.25 × N) in samples was determined by the combustion method (AOAC 2000; method 990.03) using a LECO FP-528 N (Leco Corp., St. Joseph, MI). Oil (as ether extract) in diets was extracted with diethyl ether by the ether extraction method (AOAC 2000; method 945.16) using a Soxtec system (Foss Ltd., Warrington, UK). The gross energy (GE) value of feed and excreta samples was determined in a bomb calorimeter (model 6200; Parr Instrument Co., Moline, IL) with benzoic acid used as the standard. Titanium in feed and excreta was determined as described in a previous study (Short et al., 1996).

The glutathione peroxidase assay in blood was performed using a Ransel GSH-Px kit (Randox Laboratories Ltd., UK) that employs the method based on that of Paglia and Valentine (1967). Total antioxidant status (TAS) determined in the blood serum was determined using a Randox kit, following manufacturer’s recommendations (Randox Laboratories Ltd., UK). The heterophil/lymphocyte (H:L) ratio in blood was determined as described by Müller et al. (2011). The pack cell volume (PCV) test, also called the haematocrit test, was also determined (Fedde and Wideman, 1996).

The vitamin E content in diets and livers was determined using an HPLC system as previously described (Karadas et al., 2010, 2014).

The relative empty weights of GIT segments, including spleen and heart, of each bird were determined as previously described (Abdulla et al. 2017; Pirgozliev et al. 2019a). The collected jejunal samples were stored for two weeks in 10 % neutral buffered formalin, then were embedded in paraffin wax, sectioned at approximately 5 μm and four gut segments were fixed on each slide. Morphometric measurements were determined on 20 intact well-oriented villus–crypt units for each bird as previously described (Yovchev et al., 2019).

### 2.4. Calculations

Dietary DMR coefficient was calculated using the following equation:

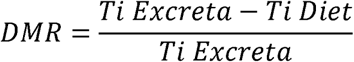

where Ti Excreta and Ti Diet are the concentrations of Ti in the excreta and diet, respectively.

Dietary NR coefficients were calculated using the following equation:

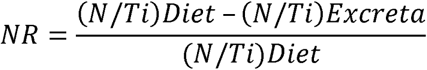

Where (N / Ti) Diet = ratio of the N to titanium in diet, and (N / Ti) Excreta = ratio of the N to titanium in excreta.

The AMEn value of the experimental diets was determined following the method of Hill and Anderson (1958).

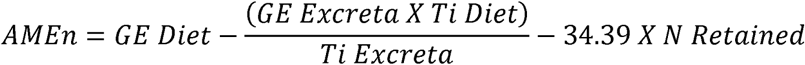

where AMEn (MJ/kg) = N-corrected apparent metabolizable energy content of the diet; GE Diet and GE Excreta (MJ/kg) = GE of the diet and excreta, respectively; Ti Diet and Ti Excreta (%) = titanium in the diet and excreta, respectively; 34.39 (MJ/kg) = energy value of uric acid; and N Retained (g/kg) is the N retained by the birds per kilogram of diet consumed.

The retained N was calculated as

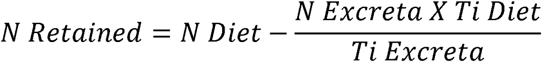

where N Diet and N Excreta (%) = N contents of the diet and excreta, respectively.

The relative development of organs was determined as follows:

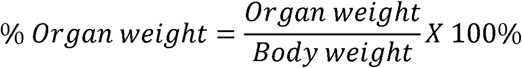

where Organ weight and Body weight are the weight of the organs and each bird, respectively.

### 2.5. Statistical Analysis

Data was analysed using Genstat (18^th^ edition) statistical software (IACR Rothamstead, Hertfordshire, UK). Comparisons among performance, diet and high temperature were performed by the general ANOVA procedure using a 2 × 2 × 2 factorial design with a split-plot structure (room). The main factors being the temperature, and the presence of dietary DHQ and vitamin E. Data was checked for normal distribution. A protected LSD test was used to separate differences in measured variable means and interaction means if differences p<0.05.

## 3. RESULTS

The analysed chemical composition of the basal diet is detailed in Table 1. The analysed protein content was lower than the calculated values, which is likely due to differences between the composition of the actual ingredients versus values used for dietary formulation. The determined values of vitamin E (µg/g) for the control and diets 2, 3 and 4 were 43.86, 54.31, 87.51 and 83.49, respectively. The supplementary DHQ product did not contain additional vitamin E.

### 3.1. Growth performance and relative organ weights

All birds were healthy throughout the study period and there was no mortality. The overall bird weight at 28d age was 988 g, with birds reared at ST at 1196 g, and birds reared at HT at 780 g (P = 0.022) (Table 2). Birds reared at HT had reduced FI and WG (P<0.05) compared to those reared at ST. Birds at HT had 35.8 % lower FI, i.e. 52 vs 81 grams daily (P = 0.020). Rearing birds at HT reduced their WG by 41 %, i.e. from 51 to 30 grams per day (P = 0.028). The FCR was not affected (P > 0.05) by diets or temperature. There was no significant effect of vitamin E or DHQ on bird production performance characteristics.

**Table 2.**
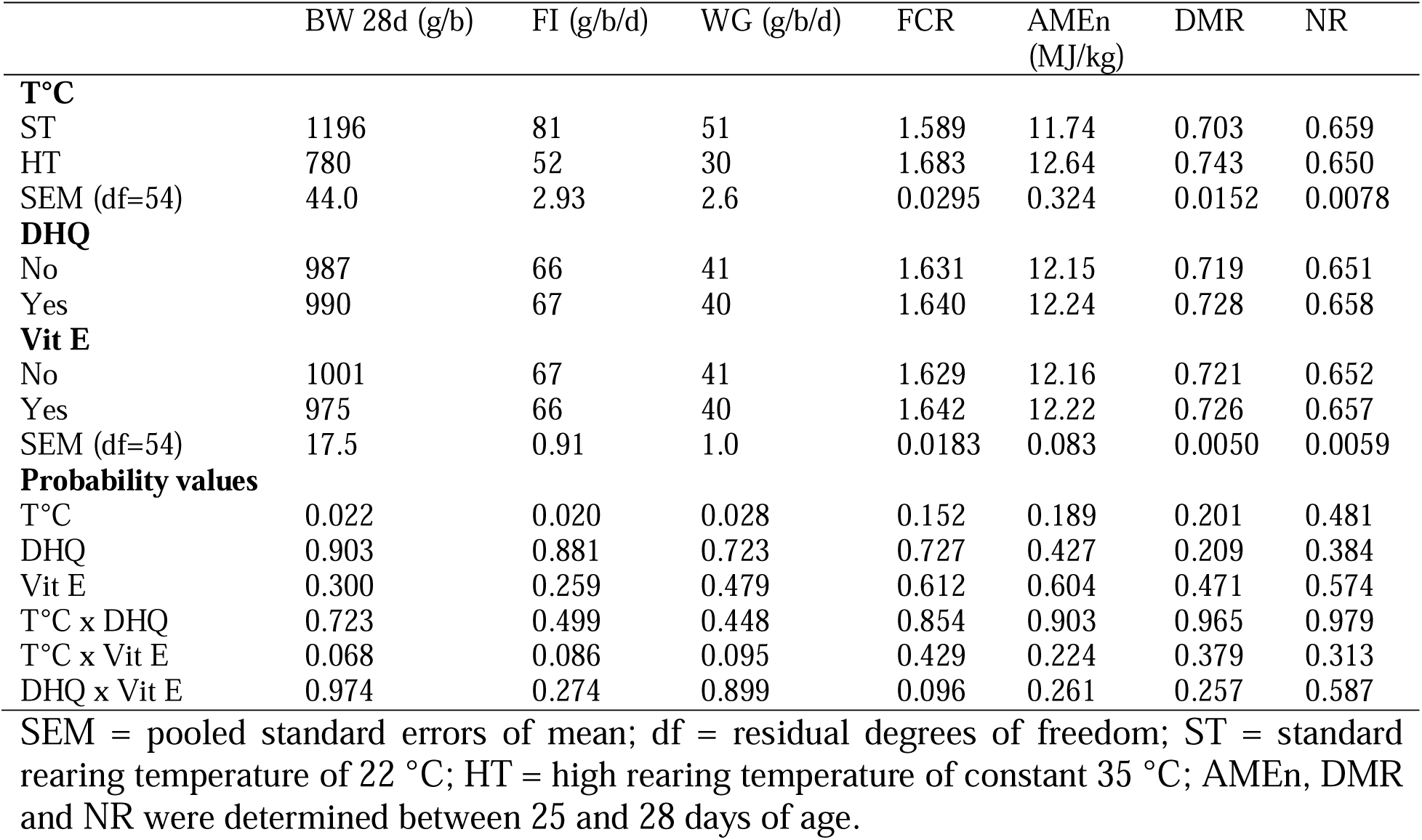
Effect of bird rearing temperature (T°C), dietary dihydroquercetin (DHQ) and vitamin E (Vit E) on final body weight (BW), daily feed intake (FI), daily weight gain (WG), feed conversion ratio (FCR), N-corrected apparent metabolisable energy (AMEn), dry matter (DMR) and nitrogen (NR) retention coefficients, when fed to broiler chickens from 7 to 28d age

The information on the development of the GIT of the birds expressed as a relative weight of the body weight is presented in Table 3. Rearing birds at HT reduced the relative weight of jejunum, liver, total GIT, spleen and heart (P < 0.05) and also tended (P = 0.091) to reduce the weight of the duodenum. Feeding DHQ significantly reduced caecal weight (P = 0.011), but increased (P = 0.002) relative heart weight. Feeding vitamin E increased the weight of liver (P = 0.011) and spleen (P = 0.009) and tended (P = 0.054) to increase the relative weight of the PG of the birds. Birds fed vitamin E reared at ST had bigger caeca (P = 0.014) compared to birds reared at HT (0.92% *vs* 0.55%), although no difference (P > 0.05) existed in birds fed diets containing no additional vitamin E (0.77% *vs* 0.65% for ST and HT respectively; data not in tables).

**Table 3.** Effect of bird rearing temperature (T°C), dietary dihydroquercetin (DHQ) and vitamin E (Vit E) on the relative organ weight expressed as the percent of body weight (BW) of gastrointestinal tract, liver, spleen and heart of 28d old broiler chickens.

|  | BW | PG | Duodenum | Pancreas | Jejunum | Ileum | Caeca | GIT | Liver | Spleen | Heart |
| --- | --- | --- | --- | --- | --- | --- | --- | --- | --- | --- | --- |
| **T°C** |  |  |  |  |  |  |  |  |  |  |  |
| ST | 1231 | 2.42 | 1.06 | 0.30 | 1.91 | 1.52 | 0.85 | 8.05 | 2.31 | 0.08 | 0.71 |
| HT | 779 | 2.48 | 0.90 | 0.31 | 1.45 | 1.27 | 0.60 | 7.01 | 1.77 | 0.05 | 0.47 |
| SEM<br>(df=54) | 45.2 | 0.057 | 0.035 | 0.012 | 0.043 | 0.105 | 0.049 | 0.110 | 0.030 | 0.001 | 0.035 |
| **DHQ** |  |  |  |  |  |  |  |  |  |  |  |
| No | 1011 | 2.41 | 0.96 | 0.31 | 1.67 | 1.43 | 0.78 | 7.57 | 2.03 | 0.07 | 0.56 |
| Yes | 1000 | 2.49 | 1.0 | 0.30 | 1.69 | 1.34 | 0.66 | 7.49 | 2.05 | 0.08 | 0.61 |
| **Vit E** |  |  |  |  |  |  |  |  |  |  |  |
| No | 996 | 2.38 | 0.95 | 0.30 | 1.70 | 1.38 | 0.71 | 7.43 | 1.97 | 0.06 | 0.60 |
| Yes | 1015 | 2.52 | 1.01 | 0.31 | 1.66 | 1.41 | 0.73 | 7.63 | 2.11 | 0.07 | 0.58 |
| SEM<br>(df=54) | 21.5 | 0.051 | 0.027 | 0.009 | 0.055 | 0.050 | 0.034 | 0.144 | 0.036 | 0.003 | 0.010 |
| **Probability values** |  |  |  |  |  |  |  |  |  |  |  |
| T°C | - | 0.552 | 0.091 | 0.654 | 0.018 | 0.231 | 0.071 | 0.022 | 0.006 | 0.001 | 0.040 |
| DHQ | - | 0.214 | 0.354 | 0.887 | 0.845 | 0.188 | 0.011 | 0.689 | 0.695 | 0.873 | 0.002 |
| Vit E | - | 0.054 | 0.165 | 0.58 | 0.570 | 0.717 | 0.678 | 0.322 | 0.011 | 0.009 | 0.257 |
| T°C x DHQ | - | 0.147 | 0.625 | 0.300 | 0.604 | 0.528 | 0.387 | 0.777 | 0.262 | 0.791 | 0.187 |
| T°C x Vit E | - | 0.627 | 0.741 | 0.376 | 0.636 | 0.900 | 0.014 | 0.750 | 0.428 | 0.556 | 0.787 |
| DHQ x Vit E | - | 0.136 | 0.683 | 0.182 | 0.923 | 0.465 | 0.823 | 0.350 | 0.797 | 0.319 | 0.264 |

SEM = pooled standard errors of mean; df = residual degrees of freedom; ST = standard rearing temperature of 22 °C; HT = high rearing temperature of constant 35 °C.

### 3.2. Dietary AMEn and nutrient availability

Dietary AMEn, DMR and NR were not influenced by supplementary DHQ, vitamin E or rearing temperature (P > 0.05).

### 3.3. Jejunal villus morphometry

The results of the jejunal villus morphometry of the chicks is presented in Table 4. There were many interactions between the studied treatments. In general, rearing birds at HT reduced VH and villus surface area without any mitigating effect from DHQ or vitamin E. It seems that feeding vitamin E and DHQ together worsened the studied villus morphometry variables reducing VH, VH:CD and villus surface area (P<0.001).

**Table 4.**
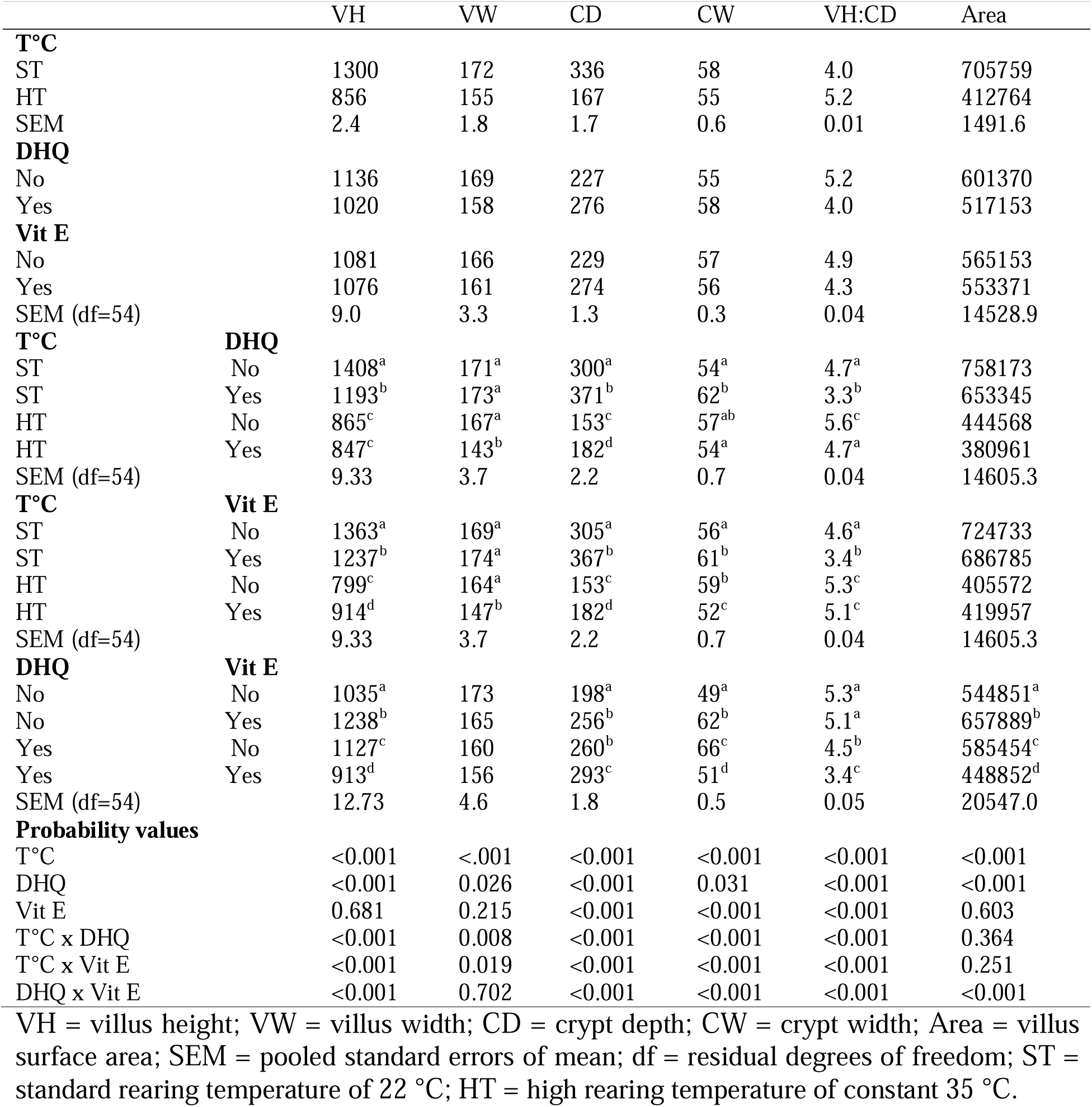
Effect of bird rearing temperature (T°C), dietary dihydroquercetin (DHQ) and vitamin E (Vit E) on the jejunal villus morphometry of 28d old broiler chickens.

### 3.4. Antioxidant status of birds

The hepatic vitamin E concentration was not affected by rearing temperature (P > 0.05) (Table 5). However, feeding DHQ or vitamin E, improved hepatic vitamin E concentration by 38.6 % and 23 %, respectively (P < 0.05). The blood GSH-Px of birds reared at HT was 17 % lower (P = 0.039) than those of birds reared at ST, i.e. 53 vs 62 u/ml RBC. However, supplementary DQH increased GSH-Px by 13 % compared to birds fed DHQ free diets (P = 0.013), i.e. 61 vs 53 u/ml RBC. Similarly, dietary DHQ improved TAS by 33.3 % (P = 0.021) compared to birds fed non-supplemented diets, i.e. 0.81 vs 0.54 mmol/L. The H:L ratio was not affected (P > 0.05) by experimental treatments. There were no diet by rearing temperature interactions (P > 0.05) for any of the studied variables in Table 5.

**Table 5.** Effect of bird rearing temperature (T°C), dietary dihydroquercetin (DHQ) and vitamin E (Vit E) on hepatic vitamin E, blood plasma glutathione peroxidase (GSH-Px), total antioxidant status (TAS), blood heterophil to lymphocyte (H:L) ratio and packed cell volume (PCV) in 28d old broiler chickens.

|  | Hepatic vitamin E<br>(µg/g) | GSH-Px<br>(u/ml RBC) | TAS<br>(mmol/l) | H:L | PCV |
| --- | --- | --- | --- | --- | --- |
| **T°C** |  |  |  |  |  |
| ST | 52 | 62 | 0.72 | 1.09 | 31.8 |
| HT | 84 | 53 | 0.63 | 1.22 | 26.0 |
| SEM (df=54) | 11.6 | 1.3 | 0.103 | 0.111 | 1.45 |
| **DHQ** |  |  |  |  |  |
| No | 57 | 53 | 0.54 | 1.11 | 29.3 |
| Yes | 79 | 61 | 0.81 | 1.20 | 28.5 |
| **Vit E** |  |  |  |  |  |
| No | 61 | 57 | 0.74 | 1.13 | 29.1 |
| Yes | 75 | 57 | 0.61 | 1.17 | 28.7 |
| SEM (df=54) | 4.8 | 2.3 | 0.078 | 0.063 | 0.57 |
| **Probability values** |  |  |  |  |  |
| T°C | 0.185 | 0.039 | 0.610 | 0.485 | 0.107 |
| DHQ | 0.002 | 0.013 | 0.021 | 0.298 | 0.315 |
| Vit E | 0.043 | 0.964 | 0.218 | 0.655 | 0.643 |
| T°C x DHQ | 0.858 | 0.094 | 0.815 | 0.470 | 0.388 |
| T°C x Vit E | 0.061 | 0.248 | 0.223 | 0.788 | 0.869 |
| DHQ x Vit E | 0.575 | 0.603 | 0.085 | 0.829 | 0.716 |

SEM = pooled standard errors of mean; df = residual degrees of freedom; ST = standard rearing temperature of 22 °C; HT = high rearing temperature of constant 35 °C.

## 4. DISCUSSION

The aim of this experiment was to evaluate the impact of dietary DHQ and vitamin E, alone and in combination, when fed to broiler chickens reared at high and standard ambient temperatures. Studying the impact of ambient temperatures is important as large variations in the temperature of poultry houses during summer months globally are increasing due to climate change (Nawab et al., 2018). There was no mortality during the entire study. The mean average weight of birds reared at the standard temperature at 28d of age was 1196 g; which is 27.5 % below the Ross 308 broiler target weight for commercial flocks. The birds were kept in small groups in research facilities, and fed mash diets, thus the reduced performance compared to large commercial flocks was expected (Pirgozliev et al., 2016; Yang et al., 2020), but was not detrimental to the study aims.

### 4.1. Growth performance and relative organ weights

In agreement with previous studies (Quinteiro-Filho et al., 2010), birds reared at a constant temperature of 35 °C responded with reduced FI and WG, although FCR was not affected by rearing temperature. The results of the relative weights of the organs measured as percentage of body weight agreed with published reports (Abdulla et al. 2016; 2017). Birds in HT group with reduced WG also had a reduced relative weight of the GIT, particularly of the small intestines. Woods et al. (2020a) also found a reduction in the relative weight of the small intestine, liver, spleen and heart in birds reared at HT. The observed reduction in the relative heart weight of birds reared at HT was further confirmed (Sonaiya et al., 1989; Yahav et al., 1999). Changes in relative organ weight may not be related to the reduced feed intake alone, since Palo et al. (1995a) found that restricted feeding only influenced absolute organ weight, not relative organ weight, and changes are transient, resulting in an improved FCR (Palo et al. 1995b). Heat stress, however, can influence hypothalamic peptides involved in appetite regulation (Song et al., 2012) and decrease feed passage rate in the GIT, further decreasing trypsin, chymotrypsin, and amylase activity (Hai et al., 2000). Chronic heat stress can reduce blood supply of the GIT due to induced peripheral vasodilation (Mckee et al., 1997), leading to a decreased size of the small intestine and absorptive capacity (Mitchell and Carlisle, 1992). High ambient temperature is therefore likely to reduce weight gain through a variety of mechanisms than the reduced feed intake alone, as noted in this study, though the effects of both factors could not be fully separated.

The enlarged hearts of the birds fed DHQ, coupled with an increase in determined GSH-Px and TAS in this study, infers that there is a potential mechanism of antioxidant protection in birds fed DHQ. However, the enlarged heart of DHQ fed birds is difficult to explain without further pathological and anatomical investigation. Korzeniowska et al. (2019) did not find differences between the relative weight of the spleen in birds fed selenium as an antioxidant. Khan et al. (2010) reported an increase in the relative weight of liver of hens with aflatoxicosis. The same authors (Khan et al., 2010) reported that a concurrent feeding of vitamin E did not ameliorate the toxic effects of aflatoxins in the hens as determined by the relative weight of the liver. Despite the liver and spleen enlargement reported in this study, no lesions and / or discolouration was observed, there was no mortality and no obvious sign of clinical disease. As previously discussed, the pathology was not determined in this study. Thus, an association cannot be made between the increase of organs size and clinical disease in this study.

The lack of response in growth performance variables to DHQ in this study is in accordance with previous research (Balev et al., 2015; Pirgozliev et al., 2019a), and is contradictory with the hypothesis that DHQ improves performance of birds reared under stress, i.e. during hot summer time (Fomichev et al., 2017). Published results on the effect of vitamin E on broiler growth performance are inconsistent as the use of vitamin E: improved performance of broilers (Guo et al., 2003) and quails (Sahin and Kucuk 2001); did not influence growth performance of broilers (Goñí et al., 2007; Niu et al., 2009; Pompeu et al., 2018) and has even reduced performance (Bölükbaşi et al., 2006). It would seem that the lack of response is prevalent in the literature and agrees with our findings in this study. However, in the reported study, the determined vitamin E in the control diet was 43.86 µg/g (65.5 IU), which is similar to the levels of dietary vitamin E recommended by the breeder (Aviagen Ltd, Edinburgh, UK) of 65 IU for this age of Ross 308. This suggests a potential explanation for the lack of response observed in growth performance variables in this and other similar studies.

### 4.2. Dietary AMEn and nutrient retention

Despite the reduction in feed intake and changes in GIT segment weights and villus morphometry, the results for AMEn and nutrient retention coefficients in the reported study were not influenced by rearing temperature or dietary antioxidants. Published data on the impact of high ambient temperature on dietary ME and nutrient digestibility are not consistent. Bonnet et al. (1997) reported a reduction in ME and nutrient digestibility values in birds reared at 35 °C, although Woods et al. (2020a) did not observe differences when studying the same variables at the same temperature. De Souza et al. (2016), Attia et al. (2018) and Habashy et al. (2017), reported an increase in nutrient digestibility in birds reared at high temperatures, while Koelkebeck et al. (1998) did not find an impact of rearing temperature on overall amino acid digestibility in laying hens. The differences may be attributed to different age, breed and type of production of the experimental birds, different dietary formulations, exposure to different temperatures for different lengths of time, ambient humidity and rearing conditions. Hai et al. (2000) reported that birds reared at a high temperature had decreased activity of trypsin, chymotrypsin and amylase, and suppressed ability to expel digesta from the crop or small intestine. Reduction in pancreatic enzyme production is usually associated with an increase in the size / weight of the pancreas in order to compensate (Kubena et al., 1983; Abdulla et al., 2016). The relative weight of the pancreas in this report was not affected by rearing temperature. Thus, suggesting that the reduced release of digesta from the crop to small intestine in HT reared birds may lead to a proportional reduction in the release of pancreatic enzymes, providing the same available energy and nutrient retention coefficients. Limited studies have reported comparisons in ME and nutrient availability in antioxidant supplemented diets. In agreement with previous reports (Goñí et al., 2007; Pirgozliev et al., 2019b), no differences were found between the broilers fed control, vitamin E and DHQ with regard to ME and nutrient retention coefficients. Studies with other antioxidants, i.e. dietary selenium, also did not detect differences in ME and nutrient retention coefficients (Choct et al., 2004; Woods et al., 2020b). As ME is a measurement of the available energy in carbohydrates, fats and proteins, it is expected that dietary antioxidants would not greatly impact the ME status.

### 4.3. Jejunal villus morphometry

The results of the villus measurements were in the expected range for birds at this age and reared under similar conditions (Santos et al., 2015; Pirgozliev et al., 2019b). Studying histo- morphometric changes in the intestines of broilers during heat stress, Santos et al. (2015) indicated that the duodenum and jejunum showed more damage than the ileum. In agreement with the reported study, Santos et al. (2015) also found that when compared with morphologically normal jejunal villi, the villi of birds reared at HT had decreased height and surface area. The increase in the number and size of the intestinal villi increases the absorption surface per unit of intestinal area (Yovchev et al., 2019), thus, HT reduces the absorptive capacity of the small intestine. However, villus morphometry does not always correlate with detectable differences in bird growth (Pirgozliev et al., 2010).

### 4.4. Antioxidant status of birds

In many cases, the antioxidant status in birds is determined by measuring TAS and GSH-Px activity (Krawczuk-Rybak et al., 2012; Mazur-Kuśnirek et al., 2019). The antioxidant enzyme system, including GSH-Px and TAS, works in concert with free radical scavengers to quench ROS and to protect cells from oxidative damage (Weiss, 1986). The balance between the production of free radicals and the antioxidant system could be disturbed by heat stress in chickens (Lin et al., 2006). As temperature increases, oxidative stress would be expected to increase and the animal’s overall GSH-Px and TAS would be expected to decrease (Ma et al., 2014; Huang et al., 2015; Sarica et al., 2017). In agreement with these reports, exposing birds to HT in this study decreased the overall GSH-Px and tended to decrease TAS.

Dietary DHQ increased GSH-Px, TAS and hepatic vitamin E which further supports the view that flavonoids can protect animal cells against oxidative stress, an action attributed to their antioxidant properties (Chen and Deuster, 2009). Supplementary vitamin E improved hepatic vitamin E, but did not affect blood antioxidant markers in this study. Feeding vitamin E with an organic source of selenium to broilers, Choct and Naylor (2004) found changes in blood GSH-Px, but not in the growth performance of the birds. Feeding dietary vitamin E at 200 mg/kg, Mazur-Kuśnirek et al. (2019) observed an improved TAS which was coupled with a higher percentage content of breast muscle. It is known that inadequate vitamin E status lowers corticosteroid synthesis and thus reduces the animal’s ability to cope with stress (Choct and Naylor, 2004)). However, the control diet of Mazur-Kuśnirek et al. (2019) showed a very low dosage of vitamin E, 10 mg/kg *vs* 50 mg/kg requirement, which may explain the observed responses. Although it was not supported with a growth response, birds with increased blood plasma and hepatic antioxidant contents may be able to cope better with the challenges when reared under standard industry conditions.

The intent was to apply the H/L ratio method to measure oxidative stress based on established principles (Maxwell and Robertson, 1998). Heat stress alters homeostasis by affecting the adrenal-corticoid axis and the resulting changes in hormone levels may change the numbers of lymphocyte and heterophil, thus changing the H/L. This method has however been criticised for not providing an adequate indication of stress alone (Müller et al., 2011; Cotter, 2015; Lentfer et al., 2015), which agrees with our results.

Broilers selected for an improved feed conversion ratio, e.g. Ross and Cobb strains, were shown to have more difficulty adapting to changes in their environment, such as extreme temperatures (that may lead to dehydration), high humidity and dietary constituents, than in less selected birds (Scheele et al., 1991). The increased PCV in modern broiler strains is associated with an increase in blood viscosity, pulmonary arterial hypertension, ascites and death (Odom, 1993; Fedde and Wideman, 1996). The PCV values in the reported study were in the expected range (Fedde and Wideman, 1996; Hasan et al., 2015). In agreement with our results, Hasan et al. (2015) also did not find differences in PCV in a study with Cobb 500, despite high rearing temperatures (22 °C *vs* 35 °C). Altan et al. (2000) further supported our results and did not report differences in PCV between birds reared at 22 °C vs 39 °C. Interestingly, low ambient temperatures can cause rapid and significant rises in PCV (Shlosberg et al., 1992) although dietary type is unlikely to affect PCV (Shlosberg et al., 1992). It seems that PCV can be changed when birds are exposed to extreme stressors, thus the lack of differences in PCV between birds fed DHQ or vitamin E was not a surprise.

Direct comparisons between studies using DHQ are difficult because there is no consistency in dietary concentrations (Pirgozliev et al., 2019a). In the reported study, DHQ was added at 0.5 g per kg feed. On average birds were consuming approximately, 67 g feed per day, and their average daily weight gain was approximately 41 g. Thus, the average daily consumption of DHQ was 0.03 g per bird, or 0.73 g per kilogram daily growth. The lack of adverse effects on animals fed relatively high dietary DHQ concentrations in the reported and in previous studies (Booth and DeEds, 1958; Pirgozliev et al., 2019a) gives an opportunity for further research, including various dietary DHQ concentrations. Studying the potential interactions between DHQ and exogenous enzymes, or comparing DHQ of different purities may also be of interest.

The mode of action of flavonoids is usually associated with their antioxidant properties (Andriantsitohaina et al. 2012), but flavonoids do not behave the same way in vitro and in vivo (Veskoukis et al. 2012). In the present study, birds fed DHQ or vitamin E had no interaction with rearing temperatures. Thus, the antioxidant properties of DHQ and vitamin E did not benefit the overall growth performance variables of birds reared at high ambient temperature. It should be noted, however, that in the reported study, the determined level of dietary vitamin E in the control diet was close to the daily recommendations of the breeder.

## 5. CONCLUSIONS

Rearing birds at a high ambient temperature reduced daily feed intake and weight gain but did not affect the efficiency of feed utilisation. Feeding DHQ or vitamin E improved various aspects of antioxidant status of the birds, although it did not affect growth performance, energy or nutrient availability. There were no observed interactions between dietary antioxidants and rearing temperature in the variables studied. At present it is unclear if feeding antioxidants (in particular DHQ) at different levels, using different dietary formulations and rearing birds under a range of environmental conditions may be effective at enhancing production performance and bird health in hot ambient climates.

## ACKNOWLEDGEMENTS

Special thanks to Richard James and Rose Crocker of the National Institute of Poultry Husbandry (Harper Adams University) for their technical support in conducting the study.

## DATA AVAILABILITY STATEMENT

The data that support the findings of this study are available from the corresponding author, upon reasonable request, subject to restrictions and conditions.

## ETHICS STATEMENT

The authors confirm that they have followed all appropriate EU and UK standards and regulations for the protection of animals used for scientific purposes. All mandatory laboratory health and safety procedures have been complied with in the course of conducting this experimental work. This manuscript complies with the ARRIVE guidelines (Kilkenny et al., 2010).

